# Disrupted Hippocampal-Prefrontal Networks in a Rat Model of Fragile X Syndrome: A Study Linking Neural Dynamics to Autism-Like Behavioral Impairments

**DOI:** 10.1101/2024.10.15.617900

**Authors:** Mohamed Ouardouz, Patrick Jasinski, Mohamed Khalife, J Matthew Mahoney, Amanda E Hernan, Rod C Scott

## Abstract

Fragile X Syndrome (FXS) is associated with autism spectrum disorder (ASD) symptoms that are associated with cognitive, learning, and behavioral challenges. We investigated how known molecular disruptions in the Fmr1 knockout (FMR-KO) rat model of FXS negatively impact hippocampal-prefrontal cortex (H-PFC) neural network activity and consequent behavior.

**Methods:** FMR-KO and control rats underwent a battery of behavioral tests assessing sociability, memory, and anxiety. Single-unit electrophysiology recordings were then conducted to measure patterns of neural activity in H-PFC circuit. Advanced mathematical models were used to characterize the patterns that were then compared between groups using generalized linear mixed models.

**Results:** FMR-KO rats demonstrated significant behavioral deficits in sociability, spatial learning, and anxiety, aligning with symptoms of ASD. At the neural level, these rats exhibited abnormal firing patterns in the H-PFC circuit that is critical for learning, memory, and social behavior. The neural networks in FMR-KO rats were also less densely connected and more fragmented, particularly in hippocampal-PFC correlated firing. These findings suggest that disruptions in neural network dynamics underlie the observed behavioral impairments in FMR-KO rats.

**Conclusion:** FMR-KO significantly disrupts several characteristics of action potential firing in the H-PFC network, leading to deficits in social behavior, memory, and anxiety, as seen in FXS. This disruption is characterized by less organized and less resilient hippocampal-PFC networks. These findings suggest that therapeutic strategies aimed at normalizing neural dynamics, such as with brain stimulation, could potentially improve behavior and cognitive functions in autistic individuals.

**HIGHLIGHTS:**

- Fragile X Syndrome is associated with autism, cognitive challenges and anxiety
- The loss of Fmr1 protein disrupts processes involved in building neural networks
- The consequence is abnormal neural dynamics in hippocampal-prefrontal cortex networks
- Normalization of dynamics could improve outcomes in FXS and ASD

## INTRODUCTION

Fragile X Syndrome (FXS) is a common monogenic cause of autism spectrum disorder (ASD)^1,2^ commonly associated with learning, cognitive and behavioral concerns that may negatively impact quality of life. Characterization of pathophysiological mechanisms responsible for the phenotype could identify novel therapeutic targets. To date, extensive evaluation at the cell biology level^3,4^ has not resulted in markedly effective therapies. We therefore shift focus to neural network dynamics, specifically action potential firing patterns, as abnormal firing patterns could ultimately represent a therapeutic target for brain stimulation strategies. The ASD phenotype in FXS implicates hippocampal-prefrontal cortex (H-PFC) neural circuit dysfunction^5,6^ and therefore we evaluated H-PFC dynamics, at the level of action potential firing patterns, in a rat model of FXS.

FXS is a disorder in which the genetic perturbation and cellular effects are known. It results from a mutation in which a DNA segment, known as the CGG triplet repeat, is expanded within the FMR1 gene. The expanded CGG triplets are hypermethylated leading to decreased expression of the *FMR1* gene with consequent absence of the FMR1 protein^7–9^. This impacts the nuclear export^7,10,11^, cytoplasmic transport and translational control of >1000 mRNAs that regulate protein synthesis, thereby modifying synaptogenesis, maturation of dendritic spines, cell-cell communication, cytoskeletal and microtubular structure, and metabolic regulators^11–20^. Many of the abnormal molecular and cellular processes identified interact to disrupt the normal maturation and structure of neural networks^21–23^ that could result in the emergence of abnormal neural network dynamics.

To define abnormal action potential firing patterns that are related to the cognitive and behavioral issues identified in FXS we studied the H-PFC circuit in FMR-KO rats. We show that FMR-KO rats have behaviors consistent with ASD and this is associated with abnormal H-PFC action potential firing patterns. This sets the stage for subsequent experiments that attempt to normalize neural dynamics to evaluate whether this improves cognitive and behavioral outcomes in FXS.

## MATERIALS AND METHODS

### Animals

All animal work was approved by the Nemours Foundation Institutional Animal Care and Use Committee in accordance with the National Institutes of Health Guide for the Care and Use of Laboratory Animals. FMR-KO and control rats were from Envigo Global service Inc Indianapolis, IN. FMR-KO rats were created at SAGE Labs in St louis, MO. All animals were housed in the AAALAC-accredited Nemours Children’s Hospital Life Science Center. Animals are single housed at a standard 12-hour light dark cycle, and water and food were provided *ad libitum*.

### Behavioral experiments

#### Social interaction

A three-chamber apparatus with a central empty chamber and 2 lateral chambers with a cage where a rat can fit is used. A rat is placed in the empty apparatus for 10 minutes to explore and then the doors are closed between the chambers. The test animal is put in the central chamber for 3 minutes while a new rat is put under a cage in one lateral chamber and an inanimate object is placed in the other. The doors are opened for 10 minutes, and the time spent interacting with the rat in the cage, or the inanimate object, is measured. The doors were closed again for three minutes with the test rat in the central chamber. A second rat is put under the cage in the lateral chamber previously housing the inanimate object. The doors are reopened for 10 minutes, and the time spent by the test rat interacting with the new and familiar rat in the lateral chambers was measured.

#### Barnes maze

The apparatus is an elevated circular platform (122 cm diameter) with circular holes (10 cm diameter) placed at the edge of the platform. A small and dark recessed chamber is placed under one of the holes and it is not visible to the animal. Pretraining: Extra-maze clues are placed in a constant position during the test. The animal is placed at the center of the maze and kept in the dark using a box for 10 seconds and then the cover is removed, and the animal is guided gently to the dark recessed chamber and allowed to stay for 1 minute. Test: the animal is placed under a box in the center of the maze for 10 seconds and then allowed to search for the hole with the dark recessed chamber. The process is repeated twice at 1-hour intervals for 5 days. The test is videotaped for post experiment analysis. The time spent to escape into the dark recessed chamber was measured. If the animal cannot find the correct hole after 3 minutes, it is gently guided to it.

#### Light Box/Dark Box

This is a two-compartment paradigm that tests for anxiety. There is a large light compartment (2/3 of the box is bright and open) and a small dark compartment (1/3 of the box). The two compartments are connected by a door that allows the animal to explore freely within the two compartments. The time spent in each compartment is recorded in addition to the number of times the animal put his head outside the dark compartment and the number of times the animal enters the lit compartment immediately following a nose poke. Animals with anxiety are less likely to spend time in, and are less likely to enter, the lit area of the box.

#### Electrophysiology recordings

Recordings were carried out as previously reported. At P45 rats were anesthetized with isoflurane (2–3% in oxygen) and placed in a stereotaxic frame (Kopf Instruments, Tujunga, CA). Custom-built electrodes were implanted using atlas coordinates into dorsal CA1 of hippocampus and medial PFC (mPFC)^24,25^. Four drivable tetrodes were placed in the hippocampus (coordinates: −3.2 mm A/P, +2.2 mm M/L, and 1.7 mm D/V) and two drivable tetrodes in the mPFC (coordinates: +2.5 mm A/P, +0.5 mm M/L, −2.5 mm D/V).

Tetrode assemblies were advanced until CA1 and PFC single-unit activity was detectable. The signal from the electrodes was preamplified at the rat’s head and transmitted to Neuralynx recording system (Neuralynx, Bozeman, MT). Putative single-unit firing was identified by clustering action potentials from the filtered (500–9000 Hz) signal and thresholded (*>*3× root mean square (RMS) noise) using Neuralynx Spike Sort 3D (Neuralynx, Bozeman, MT). Clusters were evaluated for quality using L-ratio and isolation distance in Spike Sort3D.

##### Generation of Post-Spike Filters (PSF)

A post-spike filter is a mathematical function that is proportional to the probability density that a single unit will fire an action potential (“spike”) as a function of time after it has fired a previous spike^26^. It is obtained from the spike train observed from each individual single unit by finding the probability density that maximizes the likelihood that such a spike train would occur. Guided by the knowledge that the refractory period of a neuron is roughly 2 ms we used a bin size of 1 ms to construct a binned spike train that has no more than 1 spike per bin. These data are modeled as a Poisson process from which the probability density is obtained. The probability density is written in terms of a set of basis functions and the coefficients of these basis functions are then found by numerically maximizing this expression. For the basis functions we used the raised cosine bumps of Pillow and collaborators^27^. To increase goodness of fit, penalized regression is used in which a hyperparameter is added to the (log) likelihood to penalize overfitting, and the maximization is repeated for different values of this hyperparameter to find the optimal solution between fitting the data and minimizing the penalty. For the penalized regression, we followed Pillow and collaborators^27^ by adding a term proportional to the sum of the squares of the coefficients to the log likelihood. The resulting expression is maximized for various values of the hyperparameter using Matlab on a Windows-based personal computer, to find the optimal value for fitting the data. PSFs were made to a maximum time bin of 662 ms, resulting in a 662-component vector for each post-spike filter.

##### Generation of Coupling Filters

The coupling filter of a single unit A influencing the firing of another single unit B is the probability density that B fires a spike as a function of time after A fires a spike. It can be found from the data in a very similar way as for the post-spike filter, but here the spike trains of both single units are used. Coupling filters are generated for the entire ensemble of single units from a recording session all at once. This involves adding another set of basis vectors for each probability density associated with each of the other single units, and another set of coefficients to be found, greatly increasing the cost in computational resources for the maximization problem. To do this we used a 32-core cluster running Octave instead of a PC running Matlab. Similarly, to the post-spike filters, these coupling filters were made to a maximum time bin of 662 ms for forward time only.

##### Principal Component Analysis of Post-Spike and Coupling Filters

After all the filters were made, they were normalized so that the area under each filter was equal to 1. The PSFs were grouped into two pools based on whether the single unit was recorded from the PFC or hippocampus (HC). The coupling filters were grouped into 4 pools based on whether the target and source neurons were (1) both from the HC, (2) both from the PFC, (3) target from the HC and source from the PFC, or (4) target from the PFC and source from the HC. Using Matlab, uncentered Principal Component Analysis was then performed on each pool separately in the 662-dimensional space of the components of the filters, where the number of observations corresponds to the number of filters in that pool. The first two principal components were saved and plotted, and for each filter, the first two scores were saved for further analysis. Within each pool, the PC scores were further subdivided into two groups based on whether the filter came from a control or an FMR-KO animal. The mean normalized filter for each group was also plotted for comparison, and the individual filters were plotted side by side in a heat map, sorted by PC1 and PC2 scores, respectively.

##### Similarity Analysis of Post-Spike Filters and Coupling Filters

Within each ensemble of single units from each recording session, the similarity of each filter to every other filter was quantified by first normalizing and then taking the dot product of each filter to every other filter. This was done for both post-spike and coupling filters. For each pair of normalized filters, the result will be a number between 1 for filters which are equal component by component, and −1 for filters which are the negatives of each other. We constructed graphs from PSFs where a node was a neuron’s PSF and the edge weight between nodes was the dot product. We then used a graph neural network (GNN) to characterize the normalized weighted degree and the clustering coefficient of the constructed graphs. For coupling filters, we directly compared the similarities between groups and between the regions from which the coupling filter was generated.

##### Pairwise analyses

In this analysis we binarized pairs of neurons into those with significant comodulation and those pairs that were not significantly comodulated. For each pair of neurons in a recording session, we computed the cross-correlogram from a time lag of −662 ms to a time lag of 662 ms. To determine whether the cross-correlogram showed significant coupling between each pair of neurons, we compared the sum of all values in the cross-correlogram to a statistical ensemble of 100 randomized cross-correlograms made by shifting the raster of one cluster relative to the other. This shift was made by multiplying each time stamp of one raster by some random fraction of the total time encompassing both rasters, cutting the section of the shifted raster whose time stamps were now out of the range of this total time, and appending this section to the beginning of the shifted raster. In this way the rasters are randomized with respect to one another, while preserving the statistics (mean firing rate, mean, median and standard deviation of the interspike interval) and autocorrelogram of each raster respectively. For each shifted raster a new cross-correlogram was made for a total of 100 randomized cross-correlograms. In this statistical ensemble we found that the sum of the cross-correlogram, defined as the sum of all the values of the cross-correlogram from −662 to 662 ms, followed an approximately normal distribution. If the sum of the actual cross-correlogram of the two neurons were above the 95% confidence interval around the mean of the sum of the cross-correlogram in this statistical ensemble of randomized cross-correlograms, then the actual cross-correlogram was determined to be significant. This procedure was repeated for each pair of neurons in each recording session, and the percentage of significant cluster pairs was calculated for control and diseased groups where both members of the neuron pair were recorded from the HC, where both members were recorded from the PFC, and where one member was recorded from the HC while the other member was from the PFC (mixed).

#### Statistical Analyses

A variety of approaches were used, depending on the type of analysis. For the analyses of sociability each session was considered separately and differences between groups were evaluated using a Mann-Whitney U. The same test was used for the 2 analyses for light-dark box. In all other circumstances there were multiple observations per animal and therefore we used generalized linear mixed models (GLMM) to ensure appropriate distributions for the data and to account for within-animal correlations as a function of multiple observations. For the Barnes maze the multiple observations were trials over multiple days. For the electrophysiological data the multiple observations were the number of single units recorded simultaneously. On inspection, the distributions of the principal component scores and similarity scores were frequently difficult to define and therefore we chose to perform an inverse rank-based normal transformation that resulted in a normal distribution with a mean of zero and a standard deviation of one. This is a conservative approach as the magnitude of difference is lost by ranking. Binary pairwise data was analyzed with a binary logistic approach within the GLMM framework.

## RESULTS

### Behavior

We initially carried out a range of behavioral tests to characterize social, memory and anxiety phenotypes in FMR-KO rats and controls to confirm that, in our hands, the FMR-KO rats have a phenotype that is consistent with ASD.

#### FMR-KO rats display sociability deficits

A total of 11 FMR-KO rats and 9 controls carried out a sociability task. In session 1 the animals were exposed to a novel rat and an inanimate object with the expectation that normal rats would prefer interacting with the rat rather than the object (Fig 1A). A discrimination index of 1 represents the situation in which all the interaction was with the novel rat, and a value of −1 represents all the interaction being with the object. Control rats have a mean discrimination index of 0.73 ± 0.09 (mean ± 1 s.e.) and FMR-KO rats have a mean discrimination index of 0.36 ± 0.09 (p=0.01). Therefore, the control rats have a clear preference for interacting with another animal that is not observed in the FMR-KO rats, consistent with expectations in ASD.

**Figure 1.**
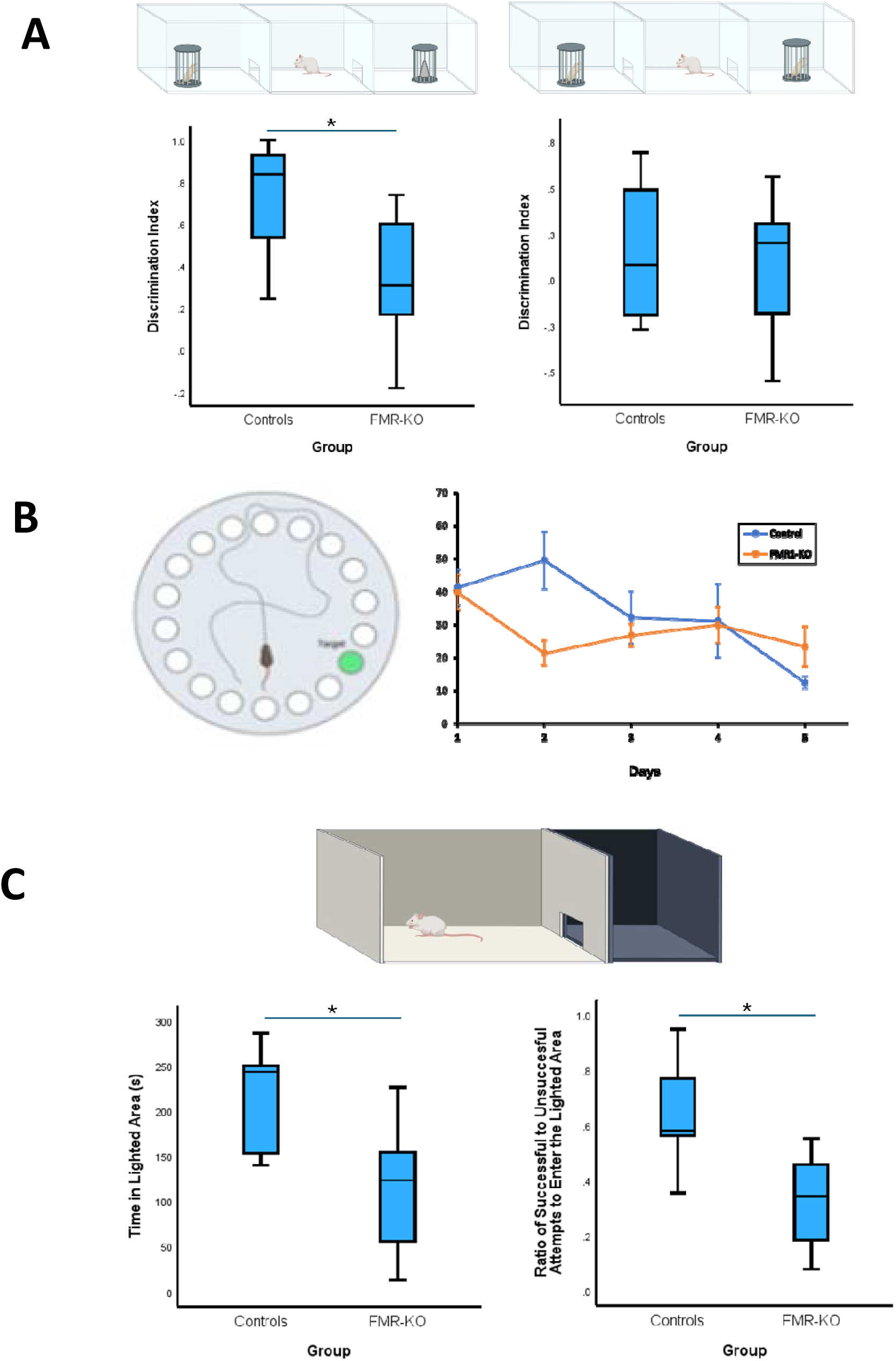
FMR-KO rats display behavioral phenotypes consistent with ASD. In a sociability task, the FMR-KO rats show significantly less preference for interacting with another rat, rather than an inanimate object, than controls. However, neither the FMR-KO or the control rats display a preference for a novel rat over a familiar one (A). The Barnes maze (B) tests spatial learning over days. The controls show an expected improvement in finding an escape hole over the course of 5 days. The FMR-KO rats have a significantly different trajectory over time such that they do not make progress after day 2. The FMR-KO rats also display an anxiety phenotype (C). They spend less time in the lit component in the light dark box and are less likely to enter the lit area after poking out their nose than control rats. (figure generated using BioRender)

In session 2 the animals were exposed to the same rat as in session 1, or a novel rat. On this occasion neither the controls nor the FMR-KO rats showed a preference for the novel rat over the familiar rat. Control rats had a discrimination index of 0.15 ± 0.13 and FMR-KO rats had a discrimination index of 0.07 ± 0.1 (p = 0.6).

#### FMR-KO rats learn a spatial task less effectively

A total of 18 rats (10 FMR-KO and 8 controls) carried out the Barnes maze task (Fig 1B). After one day of habituation, the rats had five days to learn to find a hidden escape hole in a Barnes Maze. Four large spatial cues were placed on the walls adjacent to the maze, with which the animals were expected to robustly navigate to the hidden hole. Of note, rather than this spatial strategy, animals could also visit each hole sequentially until the appropriate hole was found; this strategy is not hippocampal-dependent and usually indicates a deficit in hippocampal spatial cognition. The time it takes for the rat to find the escape hole, and how that duration changes over sessions, represents an animal’s capacity for spatial and navigational learning. The data were analyzed using generalized linear mixed models assuming a gamma distribution for latency to escape. There is a main effect of day (p = 0.006) suggesting that both groups of animals improved with increasing numbers of exposures to the environment. The mean latency to finding the escape, across both groups, on day 1 was 40.4 ± 4.2s. The latency decreased to 18.1 ± 3.7s on day 5. There is no main effect of group, averaged over days. FMR-KO animals had a mean latency to finding the escape hole, over the 5 days of 27.6 ± 2.4s, and the controls had a mean latency of 26.9 ± 2.7s (p = 0.84). However, this average measure does not provide information of how the animals are learning the task over time. On observation, it seemed that the controls have an expected strategy of using the external cues to go directly to the escape hole, and this improved over time. On the other hand, the FMR-KO rats seemed to have a strategy of testing each hole in turn until the escape hole was found. As their starting position differed each day there seemed to be more variability in time to finding the escape hole. This observation of differences in the trajectory of learning over time is strongly supported by the identification of a significant day*group interaction (p = 0.038). Pairwise analyses in the control rats showed a systematic decrease in latency from day 1 and day 2 to day 5 with significant differences as shown in Fig 1B. The FMR-KO rats had a significant reduction in latency between day 1 and day 2, but then no subsequent significant reduction in latency. On inspection of the data, the FMR-KO rats were better than control rats on day 2, but worse than the controls on day 5. Thus, the control animals showed progressive learning that was not observed in the FMR-KO rats supporting the view that the FMR-KO rats have a deficit in spatial cognition.

#### FMR-KO rats display an anxiety phenotype

Rats were placed in a light/dark box, a task based on the innate aversion of rats to brightly illuminated open areas. A total of 13 rats were used for this experiment (5 controls, 8 FMR-KO) (Fig 1C). The data were analyzed in two complementary ways. First, we evaluated the amount of time that the animals spent in the lighted area. Animals that are less anxious will tend to venture into the lighted compartment more often, spending more time there, than more anxious counterparts. Controls spent a median of 243s (IQR; 146 to 268s) compared to a median of 123s (IQR; 38-159s) in FMR-KO rats (p = 0.03). We also evaluated the proportion of attempts to enter the lighted area that resulted in going into that area, an index of activity-exploration. Control rats entered the lighted area on 58% of attempts (IQR; 46 to 86%) compared to 34% of attempts (IQR; 14 to 47%) in FMR-KO rats (p = 0.016). Thus, both analyses support the view that FMR-KO rats are more anxious than controls.

### Neural Dynamics

To understand the neural network mechanisms for these behavioral impairments, we recorded extracellular single units simultaneously from the hippocampus and the PFC, two brain regions implicated in learning and memory, anxiety, and social behavior. A total of 115 hippocampal neurons (58 in controls and 57 in FMR-KO) and 127 PFC neurons (64 in controls and 63 in FMR-KO) in 15 rats (8 control and 7 FMR-KO) were recorded. There were no differences between the groups in terms of firing rate, median interspike interval or action potential half width in either the hippocampus or the PFC.

#### Action potential firing patterns are abnormal in hippocampus and PFC in FMR-KO rats

The post-spike filters were used to evaluate temporal modulation of neuronal firing and estimate patterns of action potential firing over the millisecond (ms) timescale. The filters were dimension reduced using a principal components analysis so that PC scores could be quantified and compared between groups. The first two principal components in each brain region were evaluated.

In the hippocampus (Fig 2A), the first principal component (PC1) accounted for 57% of the variance and the second principal component (PC2) accounted for 9% of the variance. PC1 captured the bursting activity that is typical of hippocampal pyramidal cells. The neurons have a propensity of firing soon after they have previously fired, with subsequent increased propensity to fire about 120ms later; a burst firing pattern that is modulated in theta. The PC1 score reflects how close the pattern of firing of a neuron is to that canonical pattern. The mean rank normalized PC1 score in the controls was 0.22 ± 0.1 in the controls and −0.23 ± 0.1 in the FMR-KO rats (P<0.001). PC2 mostly captured an immediate increase in firing propensity followed by oscillatory behavior around a gain of zero. Most of the variance captures an increase in firing for about 10-30ms after a prior spike. The normalized mean rank PC2 score in the controls was −0.29 ± 0.09 and 0.16 ± 0.1 in the FMR-KO rats (P < 0.001). Therefore, FMR-KO hippocampal neurons have excessive early bursting behavior and less theta modulation than in controls.

**Figure 2.**
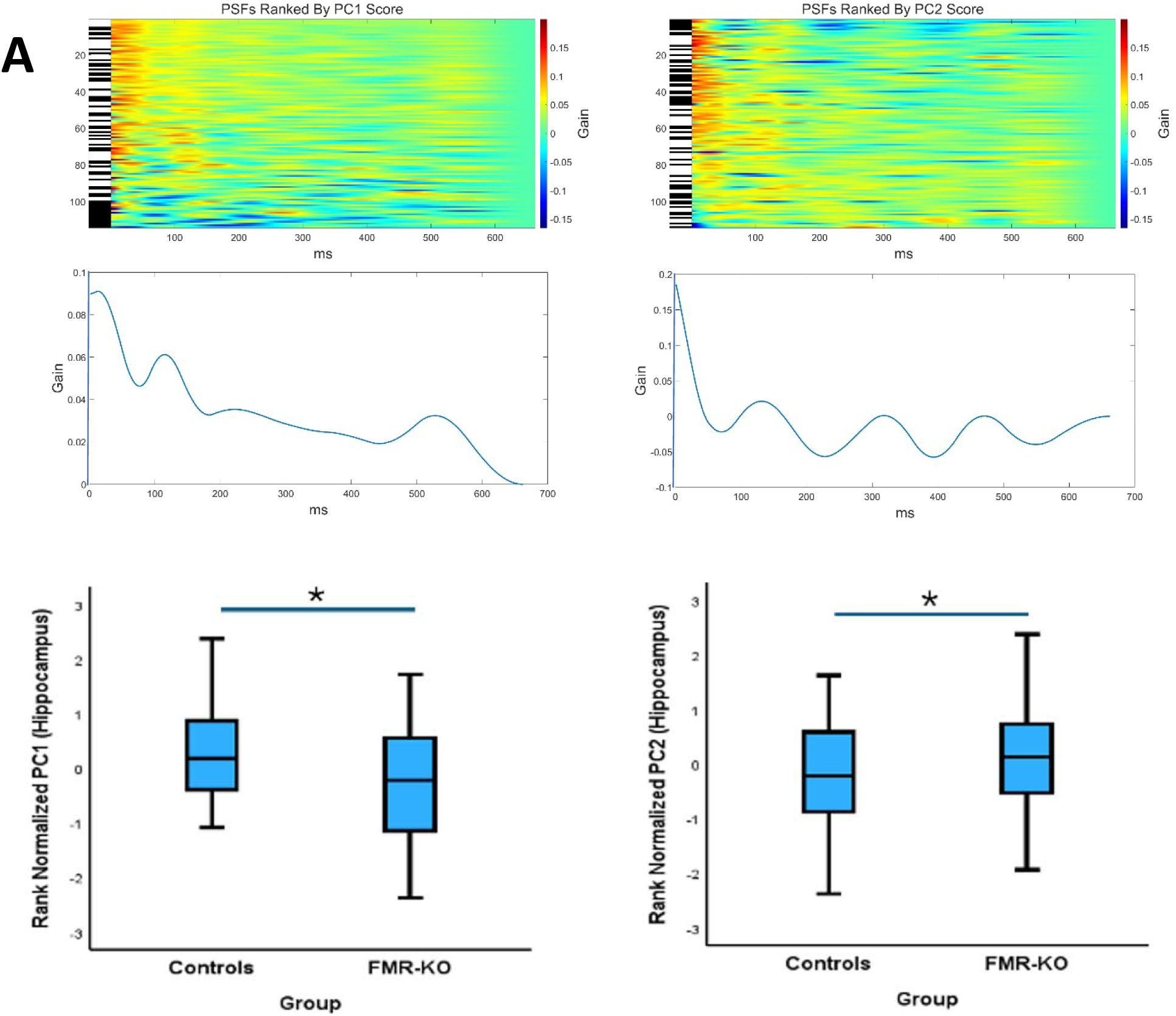
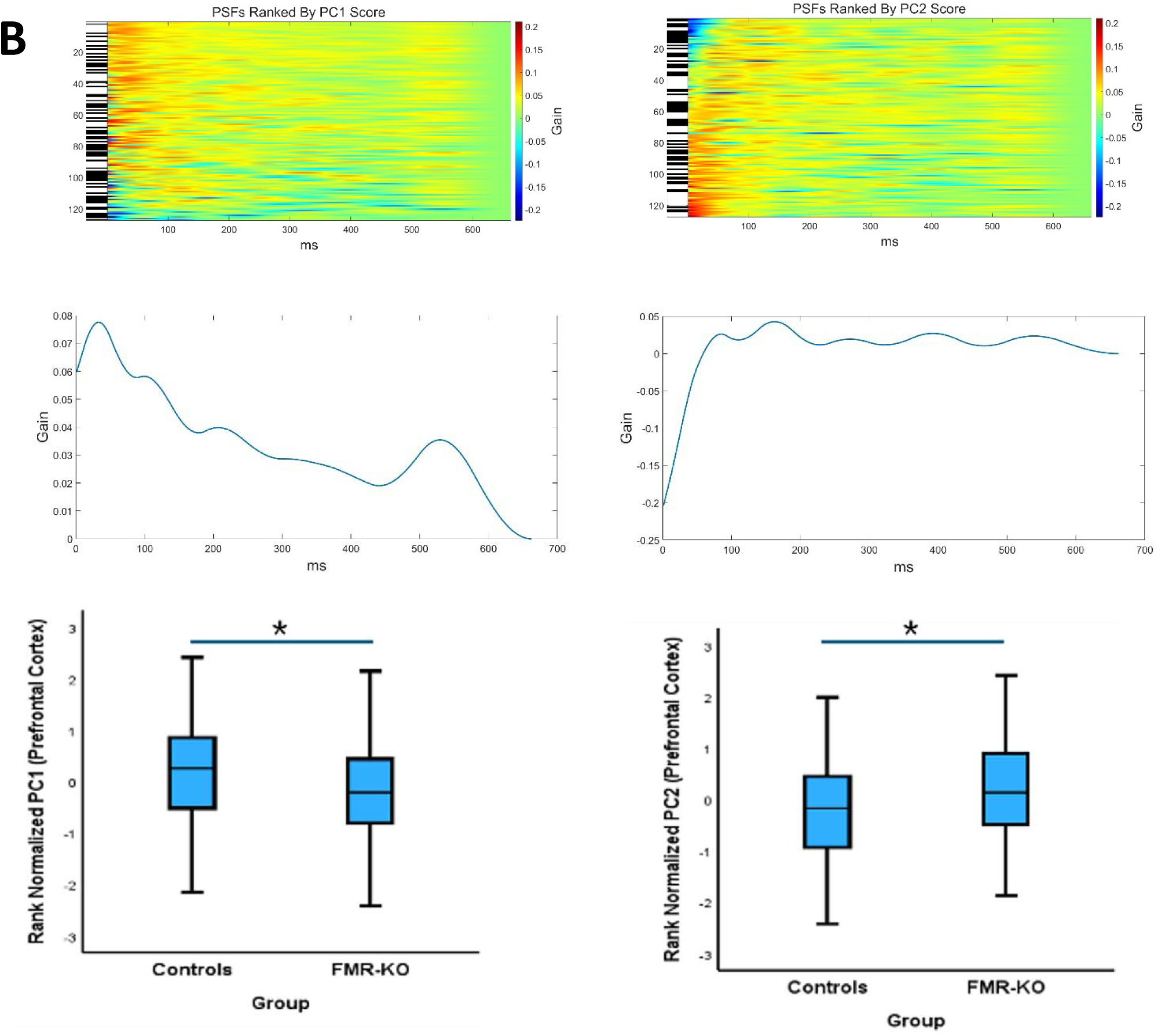
FMR-KO rats have abnormal action potential dynamics in the hippocampal-PFC circuit. A heat map of all the hippocampal post spike filters is shown in (A), and for PFC dynamics is shown in (B). The black marks to the let of the heat map show the PSFs from FMR-KO animals. The middle row shows the first 2 principal components (PC) defining the shapes that account for the majority of variance in firing. The first PC from hippocampal recordings shows the neurons have an increased propensity to fire immediately after they have just fired, followed by another increased propensity about 120ms later (theta modulation, consistent with the expectation for hippocampal neurons). The second PC from the hippocampus reveals an increased propensity to immediate firing following an initial spike with little modulation subsequently (line fluctuates around a gain of zero). In the PFC, the first PC reveals a propensity to fire about 30ms after previously firing, with a slow decay in that probability over time. The second PC is dominated by a prolonged refractory period immediately after firing. These two PCs suggest a regular firing pattern, consistent with expected behavior of PFC neurons. The box plots are the quantification of PC scores showing that FMR-KO rats have less theta modulation and increased propensity to fire immediately after firing in the hippocampus, and less robust regular firing in the PFC.

In the PFC (Fig 2B), PC1 accounted for 57% of the variance and PC2 accounted for 13% of the variance. PC1 captures a pattern with most firing occurring approximately 30ms after a previous spike, with a subsequent smooth decay in firing propensity for the next 600ms, consistent with expected behavior of PFC neurons. The mean rank normalized PC1 in the PFC is 0.19 ± 0.1 in controls and −0.20 ± 0.1 in FMR-KO rats (p=0.01). PC2 captures an immediate reduction in the propensity to fire for about 80 ms followed by a period in which firing could occur at any time. The observation that PFC neurons have the capacity to fire across the entire time captured by the PSF suggests that the PFC has capacity for flexible firing. The mean rank normalized PC2 was −0.24 ± 0.09 in the controls and 0.33 ± 0.08 in the FMR-KO rats (p = 0.001). Therefore, in FMR-KO rats the PFC neurons are less likely to fire in a regular firing pattern and are more likely to have a prolonged refractory period than in control rats.

#### Network structure based on action potential firing patterns are abnormal in FMR-KO rats

We next constructed graphs based on correlations in patterns of action potential firing across neurons, defined by the dot product of the post spike filters. Each node in the network is a neuron and the edges are weighted by the dot product between all 2 node pairs and normalized to the number of total nodes in the network. This generates a fully connected, normalized, weighted, undirected graph for each session.

The weighted degree is the sum of weights of the edges from each node and is a measure of the density of network connections (Fig 3). The weighted degrees were rank normalized to ensure that the distribution was valid. The mean weighted degree in controls was 0.42 ± 0.8 and in FMR-KO rats −0.39 ± 0.8 (P<0.001), after adjusting for whether the neuron was recorded from the hippocampus or the PFC. There was a trend for a significant group*region interaction (p=0.09). In controls the weighted degree was larger in the hippocampus than in the PFC showing that when hippocampal neurons fire, the pattern of firing is similar across neurons whereas the firing in the PFC has more variance, consistent with increased flexibility in firing patterns. This difference in pattern similarity was not observed in the FMR-KO rats.

**Figure 3.**
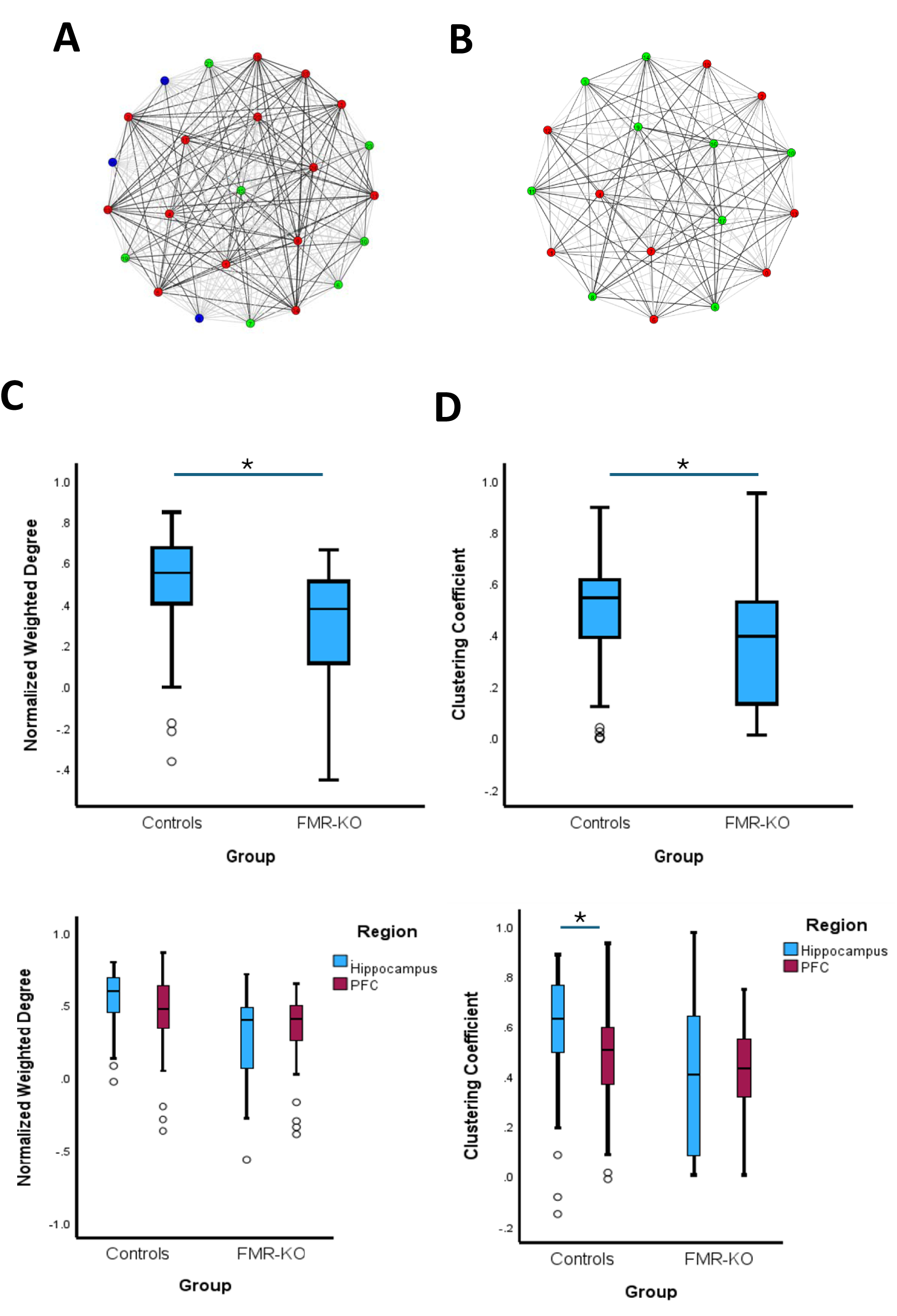
FMR-KO rats have less organized firing in the hippocampus and less flexible firing in the PFC. Example networks from a control rat (A) and and FMR-KO rat (B) are shown. The weighted degree (C) provides information on the similarities of patterns of firing between neurons in the same animal. There is less similarity in firing patterns in the FMR-KO rats when all neurons are considered. There is no difference between hippocampal similarity and PFC similarity in either the FMR-KO rats or the controls. The clustering coefficient (D) is a measure of the resiliency of the firing patterns. The FMR-KO animals have lower resiliency than controls. In this circumstance there is a difference in resiliency between the hippocampus and PFC in controls but not in FMR-KO rats.

The clustering coefficient is a measure of how connected a neuron is to its neighbors and is a measure of network resiliency. A high clustering coefficient indicates that nodes are tightly connected to form a community structure. A low clustering coefficient, on the other hand, suggests a more decentralized or random network structure. The mean rank normalized clustering coefficient in controls was 0.36 ± 0.08 and −0.32 ± 0.08 in FMR-KO animals (p<0.001, Fig 3), suggesting that control networks are more resilient than those in FMR-KO rats. On this occasion there was a significant group*region interaction term (p=0.007). In FMR-KO the mean difference in rank normalized clustering coefficient between the hippocampus and the PFC was 0.1 ± 0.16 (p=0.52) showing that there is similar, albeit low, resiliency across brain structures. In controls the mean difference in rank normalized clustering coefficient was −0.5 ± 0.16 (p=0.001), with the clustering coefficient being lower in the PFC.

#### There are fewer significant cross-correlations in FMR-KO

The above approach does not allow for direct functional relationships in time between neurons to be characterized. For each pair of neurons, we determined whether there was significant comodulation and compared the proportion of significantly comodulated pairs between groups in hippocampal-hippocampal, PFC-PFC and hippocampal-PFC pairs (Fig 4). Control animals were

**Figure 4.**
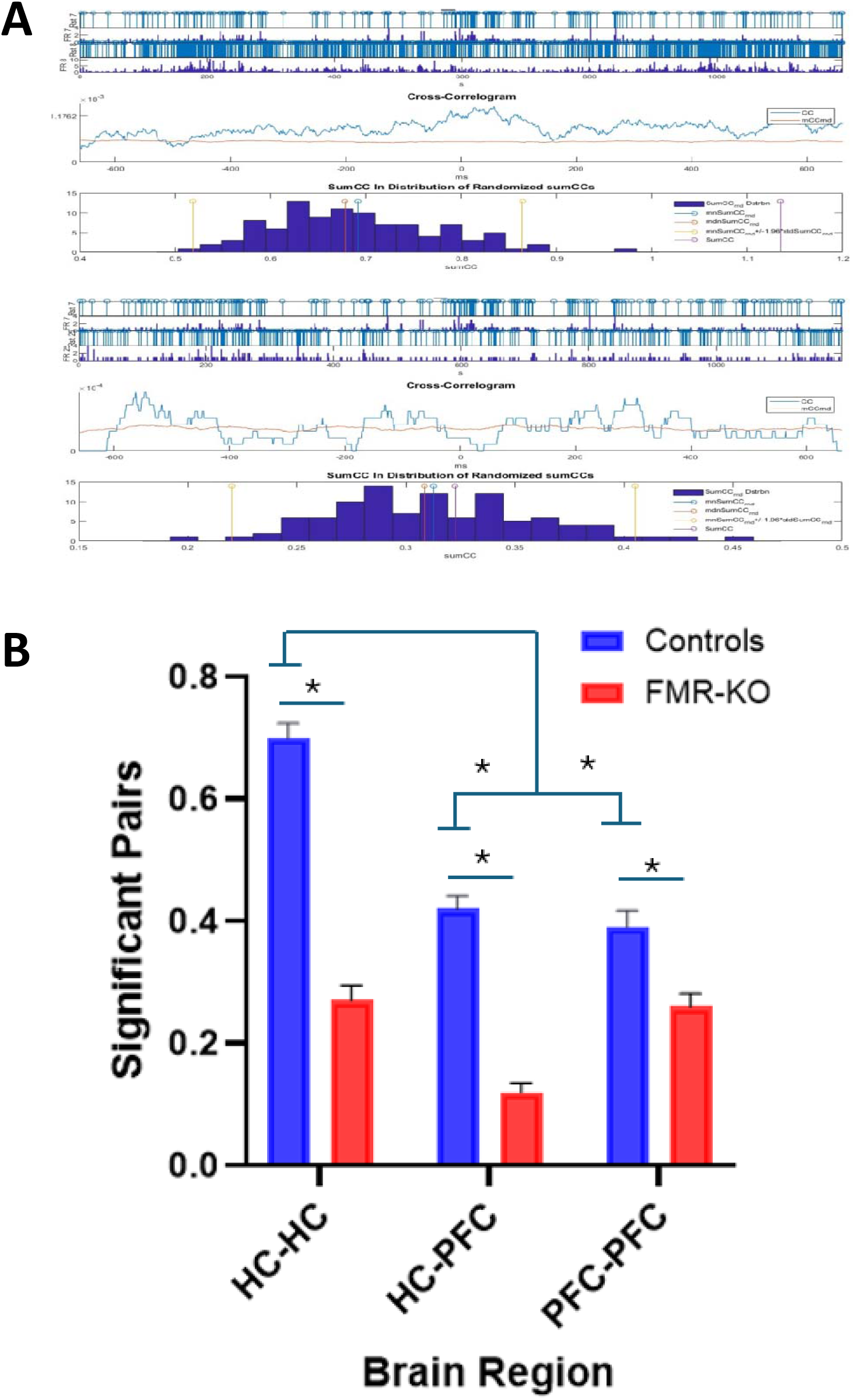
FMR-KO rats have less coordinated firing in populations of neurons. The left panel shows examples from 2 pairs of neurons, 1 pair that is comdulated (top panel in A) and1 pair that is not comodulated (bottom panel in A). Each panel shows raster plots from 2 neurons followed by the observed cross-correlogram. The sum of all the values in the actual cross-correlogram is then compared to the sum of all values from 100 randomized raster plots (histogram represents the null distribution). The mean, median and 95% confidence intervals are shown on the null distribution. In the top plot, the observed sum is very far outside the upper 95% confidence interval of the null distribution, representing comodulation. In the second example the observed sum is close to the mean of the null distribution, representing lack of comodulation. The proportion of comodulated pairs is shown for hippocampal-to-hippocampal, hippocampal to-PFC and PFC-to-PFC pairs in controls and FMR-KO rats. In all circumstances that comodulation is significantly less in FMR-KO animals. There is also a different pattern of comodulation. In controls the most comodulation was observed in the hippocampus with similar comodulation in HC-PFC and PFC-PFC pairs. In the FMR-KO animals there was similar comodulation in HC-HC and PFC-PFC pairs, with a reduction in comodulation of HC-PFC pairs.

6.2 times (95% CI; 4.5-8.7) more likely to have significant comodulation between pairs of hippocampal neurons than FMR-KO rats (p<0.001). For PFC-PFC pairs the controls were 1.8 times (1.3-2.5) times more times more likely to have significant comodulation than the FMR-KO pairs (p<0.001), and for hippocampus-PFC pairs the controls were 5.4 times (3.9-7.5) more likely to have significant comodulation than FMR-KO pairs (p<0.001). There is also a significant Group*Region interaction (p<0.001). In the controls, the hippocampal-hippocampal pairs are most likely to have significant comodulation with the proportion of comodulated pairs being similar for hippocampus-PFC and PFC-PFC. In contrast, in the FMR-KO the proportion of hippocampus-hippocampus and PFC-PFC pairs is similar (albeit lower than in controls) with a very low proportion of hippocampal-PFC pairs being comodulated.

#### Coupling filters display a large variability in firing patterns

Cross-correlations capture information about how neurons are interacting with each other in a way that is a composite of autocorrelation structure (as captured in the PSF) and any direct comodulation that is over and above the autocorrelation structure. To distinguish autocorrelation structure from the direct comodulation structure we built coupling filters (CFs). The CFs were categories into 4 groups: hippocampal to hippocampal, PFC to PFC, hippocampal to PFC, and PFC to hippocampal neurons. The first and second principal components were compared between groups in each of the above categories, however, in the HC to HC group, only 17% and 11% of the variance was captured, in the PFC to PFC group, only 16% and 11% of the variance was captured, in the HC to PFC group, only 15% and 12%, and in the PFC to HC group, only 14% and 12% of the variance was captured. This suggests that there are many different types of coupling dynamics and therefore we did not pursue this approach further.

#### Differences in similarity of CF between groups is a function of brain region

The coupling filter describes the pattern of action potential firing in a neuron, given that another neuron has just fired. The PC scores account for very little of the variance suggesting that there may be many ways in which any neuron is influenced by any other neuron. However, this does not mean that there are not similarities in co-firing patterns. To characterize the similarity in patterns of interactions between neurons we calculated the dot products between coupling filters, acknowledging that this approach will not give information on the structure of the coupling filter (Fig 5). This generated a total of 634,063 dot products in controls and 812,685 dot products in FMR-KO animals. There is no difference between the groups when all the dot products were compared (p=0.76). We then separated the data into dot products from hippocampal to hippocampal, PFC to PFC and hippocampus to PFC filters. The patterns of action potential co-firing in hippocampus-to-hippocampus pairs are more similar in controls than in FMR-KO animals (p=8.8×10^−9^). Conversely in both the PFC to PFC (p=0.0008) and hippocampus to PFC (p=3.3×10^−13^) coupling filters the patterns of firing are less similar in controls when compared to FMR-KO animals.

**Figure 5.**
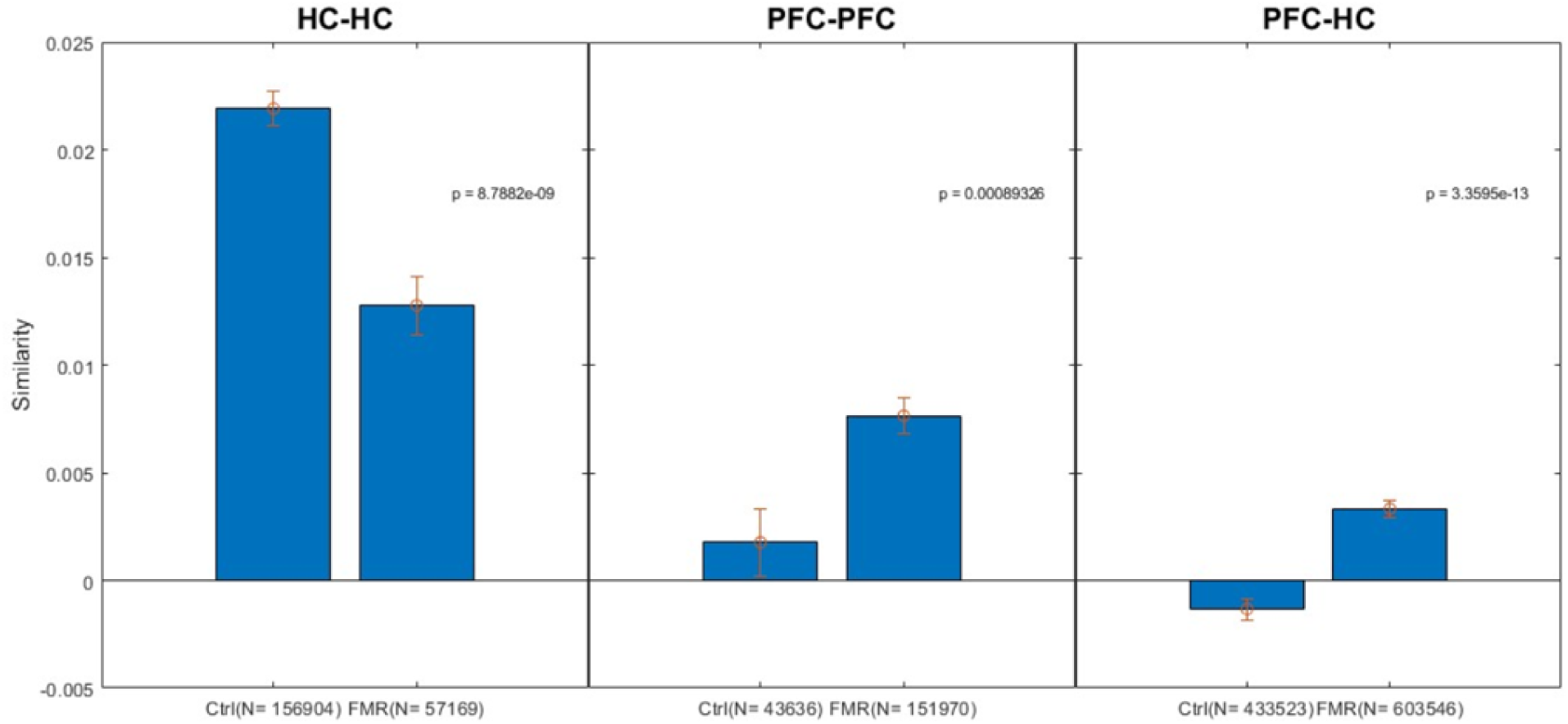
FMR-KO rats have abnormal patterns of comodulation. In controls the patterns of comodulation are more similar than that observed in FMR-KO rats. However, in hippocampal-to-PFC and in PFC-to-PFC pairs, the patterns of comodulation are less similar in controls than that observed in FMR-KO rats.

## DISCUSSION

The results of the current study confirm that loss of Fmr1 protein has major impacts on the functional architecture of action potential firing and that this is associated with important negative effects on social, memory and anxiety behaviors. These findings support the view that abnormal action potential dynamics are an important mechanism of disease pathophysiology in FXS. Therapeutic strategies such as brain stimulation targeted at recovering abnormal neural dynamics towards normal could result in improvements in cognition and behavior, thereby helping to maximize quality of life of autistic people.

The investigation of the pathophysiology of FXS has emphasized understanding the downstream impacts of loss of Fmr1 protein on a variety of interacting cell biological pathways. Many of those pathways are critical for the development and ultimate structure of neural networks that support the generation of action potentials. The formation, organization and maintenance of dendritic spines is abnormal in FMR-KO^28–32^ suggesting that inputs into the hippocampal – PFC network are inadequately integrated with resultant abnormalities in the action potential outputs of the system. The current study extends previous studies by explicitly evaluating those output firing patterns of neurons.

The evaluation of these patterns is critical as the action potential is the fundamental unit of information transfer in the brain and thus underlies all behavior. This is easy to understand with respect to patterns of action potential firing at the time of a behavior. However, it is also known that the patterns of action potential firing in one circumstance can predict behavioral performance in a different circumstance. For example, abnormalities in the default mode network^33,34^, dorsal attention network^35–37^, frontoparietal control network^38,39^ and the cingulo-opercular network^40^ all predict cognitive performance in people with Alzheimer’s disease. In experimental models, the fidelity of hippocampal CA1 place cell firing predicts performance in a maze testing spatial cognition^41,42^.This suggests that there is a baseline functional architecture of neural networks, and the integrity of that network is important for the emergence of specific action potential firing during tasks. On this basis we recorded action potential firing patterns while an animal is walking in an open field rather than when the animal was engaged in an active sociability, spatial memory, or anxiety task. We posit that the electrophysiological abnormalities identified in the current study represent an abnormal substrate that cannot generate the appropriate action potential firing patterns that are required for normal behaviors. This is an important concept as it suggests that treatments to normalize background dynamics may then allow more normal firing patterns to emerge during a range of specific tasks and is thus a potentially more generalizable approach than modifying action potential firing patterns only at the time that specific tasks are being completed. This will require validation in future studies.

Action potential firing dynamics were evaluated in several ways. The initial approach was to establish whether there are abnormalities when neuronal firing is evaluated on a neuron-by-neuron basis. The firing patterns are outputs given that neuron’s inputs. In the hippocampus, neurons typically display burst firing with bursts occurring at theta frequencies^43–46^. It is known that neurons with this behavior are also neurons that effectively encode space and events as determined by the fidelity of place cell firing^26^. There were more neurons with this canonical behavior in controls than in FMR-KO animals suggesting that control neurons have more cognitive capacity, particularly in the domain of memory, than FMR-KO neurons. In contrast, the PFC is important for executive functions such as attention, working memory, behavioral flexibility and decision making, and has been shown to be crucially involved in social behavior and anxiety. These behaviors require a network that is flexible and able to rapidly adapt to inputs. In agreement with this, spontaneous activity of neurons in the PFC is characterized by sporadic firing of weakly correlated pairs of neurons, with some neurons oscillating in delta frequency, punctuated by occasional synchronized activity^47^. In our dataset, this is manifest as a firing preference about 30ms after a neuron has just fired and a subsequent smooth decay. In addition, there is a low but uniform probability that the neuron can fire any time after a refractory period, implying sufficient flexibility in the network to allow firing as and when it is needed.

We next evaluated similarities in patterns of action potential firing within individual animals and across the hippocampus and PFC to determine whether networks built using dot products were different across groups. Each neuron is represented by a node in a graph and the dot product between nodes is the weight of an edge in a graph. We examined two standard graph parameters: the normalized weighted degree and the clustering coefficient. The weighted degree is a measure of the similarity of action potential firing patterns and is higher in the controls than the FMR-KO animals. This analysis implies that when control hippocampal neurons fire, they tend to have similar firing patterns, even if the neurons are not firing at the same time. The clustering coefficient is a measure of robustness of a network and thus provides information about resilience against random damage and is also higher in controls that in the FMR-KO animals. Thus, the loss of FMR1 protein leads to the emergence of neural networks that are unable to generate robust and similar patterns of action potential firing that support hippocampal-PFC function, and in addition are less able to tolerate any random perturbations that may occur.

There is an interesting difference between network parameters in hippocampus and PFC. In controls there is greater hippocampal firing similarity across neurons and a greater resiliency to random damage than in PFC. This suggests that in the physiological situation there is tightly organized information sent from the hippocampus and that this information is integrated into the PFC in a way that allows flexible outputs, consistent with the requirements for normal executive function. This difference is not observed in the FMR-KO rats with the major reason being that the hippocampus is less resilient to damage. This could be interpreted as poorly integrated information in the hippocampus generating inappropriate outputs that become inputs to the PFC that negatively impacts flexible firing in the PFC.

The above analyses evaluated patterns of neuronal firing independently of how other neurons were behaving. We next evaluated how neurons were behaving with respect to each other. We initially took a binary approach in which neurons were either coupled or not. We show that FMR-KO animals have far less comodulated firing both within structure (i.e. hippocampal-hippocampal and PFC-PFC pairs) and across structures (hippocampal-PFC pairs). To try and understand the dynamical patterns that link pairs of neurons we took a coupling filter approach. On observation there are an enormous number of patterns identified and this is borne out by the fact that the principal component scores only account for a very small percentage of the variance. The large number of potential patterns does not lend itself to a principal component approach as there was no specific shape that captured a lot of the variance in the data. Nevertheless, this does not exclude the possibility that there are similarities in the patterns of firing when coupling filters are correlated with each other. That analysis identified more similarity in hippocampal-hippocampal couples in controls than FMR-KO, further supporting the view that the normal hippocampus is a highly organized and integrated functional neural network. In contrast, the similarity in coupled firing in the PFC-PFC couples is lower in controls than in FMR-KO. We posit that this functional organization is appropriate for the PFC as the network receives highly organized information that is integrated in a way that has the network poised to generate short lasting interactions between neurons as the need arises. The more organized firing in the PFC of FMR-KO means that there are less opportunities for flexible functional interactions, and this is likely manifest as more rigid behaviors, consistent with ASD.

In summary, the FMR-KO rat displays several behavioral changes that are consistent with changes identified in autistic people. At the level of the neural network, the inputs to the hippocampal-PFC network are disrupted given the previously identified abnormalities at dendritic spines. This generates a network that is less organized, less integrated, and less resilient than expected in controls. These abnormalities are potential therapeutic targets for brain stimulation in which inputs are modified in a way that normalizes the background dynamical abnormalities, modifying synaptic weights as a function of activity dependent plasticity which is turn could normalize behavior. Ultimately, such targeted brain stimulation could help to maximize quality of life in autistic people.

